# How Older Adults Maintain Lateral Balance While Walking on Narrowing Paths

**DOI:** 10.1101/2024.01.04.574192

**Authors:** Meghan E. Kazanski, Joseph P. Cusumano, Jonathan B. Dingwell

## Abstract

**Background:** Older adults have difficulty maintaining side-to-side balance while navigating daily environments. Losing balance in such circumstances can lead to falls. We need to better understand how older adults adapt lateral balance to navigate environment-imposed task constraints.

**Research Question:** How do older adults adjust mediolateral balance while walking along continually-narrowing paths, and what are the stability implications of these adjustments?

**Methods:** Eighteen older (71.6±6.0 years) and twenty younger (21.7±2.6 years) healthy adults traversed 25m-long paths that gradually narrowed from 45cm to 5cm. Participants switched onto an adjacent path when they chose. We quantified participants’ lateral center-of-mass dynamics and lateral Margins of Stability (*MoS*_*L*_) as paths narrowed. We quantified lateral Probability of Instability (*PoI*_*L*_) as the cumulative probability that participants would take a laterally unstable (*MoS*_*L*_<0) step as they walked. We also extracted these outcomes where participants switched paths.

**Results:** As paths narrowed, all participants exhibited progressively smaller average *MoS*_*L*_ and increasingly larger *PoI*_*L*_. However, their *MoS*_*L*_ variability was largest at both the narrowest and widest path sections. Older adults exhibited consistently both larger average and more variable *MoS*_*L*_ across path widths. Taken into account together, these resulted in either comparable or somewhat larger *PoI*_*L*_ as paths narrowed. Older adults left the narrowing paths sooner, on average, than younger. As they did so, older adults exhibited significantly larger average and more variable *MoS*_*L*_, but somewhat smaller *PoI*_*L*_ than younger.

**Significance:** Our results directly challenge the predominant interpretation that larger average *MoS*_*L*_ as indicating “greater stability”, which we argue is *inconsistent* with the principles underlying its derivation. In contrast, analyzing step-to-step gait dynamics, together with estimating *PoI*_*L*_ allows one to properly quantify instability risk. Furthermore, the adaptive strategies uncovered using these methods suggest potential targets for future interventions to reduce falls in older adults.

## INTRODUCTION

People regularly adjust walking [1] to navigate paths constrained by fixed [2], moving [3] and/or irregular [4] features of their environments (living spaces, store aisles, sidewalks, etc.). Older adults frequently fall while navigating such contexts [5]. Narrow paths present particular challenges, limiting how much older adults can adjust foot placements to maintain mediolateral balance [6, 7]. Further, walking humans are intrinsically less stable laterally [8], and sideways falls by older adults are prevalent and injurious [9]. To prevent such falls, we must better understand how older adults maintain mediolateral balance along constrained paths.

The lateral Margin of Stability (*MoS*_*L*_) is a mechanically-grounded [10, 11] and widely used [12] metric for which larger *MoS*_*L*_, averaged over multiple steps, is often interpreted to indicate greater gait stability [12, 13]. However, this interpretation yields paradoxical findings [13]: older adults [14], persons with various gait impairments [12], or those who experience destabilizing perturbations [15, 16] often exhibit larger (supposedly *more* stable) average *MoS*_*L*_, despite clearly *de*stabilizing circumstances. Such counterintuitive findings arise because interpreting average *MoS*_*L*_ in this way is *inconsistent* with the physical principles underlying its formulation. Despite its name, *MoS*_*L*_ does not quantify “stability” in the dynamical sense of the speed with which a system recovers from perturbations [17]. Instead, true to its name, *MoS*_*L*_ quantifies the *margin* (as a linear distance) away from some *threshold* (i.e., *MoS*_*L*_ = 0) [10, 11] above which stability is expected to hold. Precisely because of this, computing the average *MoS*_*L*_ over any sequence of multiple steps is the wrong statistic to extract from such data. Indeed, people can successfully walk with a range of average *MoS*_*L*_ values and even routinely take (supposedly “unstable”) steps with *MoS*_*L*_ < 0 without falling [13]. This confusion is precisely what produces the apparent paradoxes cited above.

Here, we instead emphasize that the threshold interpretation of *MoS*_*L*_ is crucial. When a walker exhibits *MoS*_*L*_ < 0, this violates Hof’s condition for dynamic stability [10, 11] and the walker must exert additional effort to avoid a fall. The relevant statistic to compute from some sequence of steps should thus quantify the *likelihood* a walker will violate Hof’s condition [13]. We proposed the lateral Probability of Instability (*PoI*_*L*_) to directly quantify this likelihood that (on average) a given step will exhibit *MoS*_*L*_ < 0 [13]. *PoI*_*L*_ quantifies not whether a walker *will* be unstable, but that it risks *becoming* unstable on any given step [13]. *PoI*_*L*_ correctly captured people’s increased risk of losing lateral balance when destabilized, despite their exhibiting larger average *MoS*_*L*_ [13]. Here, we proposed *PoI*_*L*_ can elucidate how older adults mitigate their risks of becoming laterally unstable when walking down slowly narrowing paths.

Narrow walking paths accentuate age-related mediolateral balance impairments [18-21] and can help discern fallers from non-fallers [22]. On narrow paths, younger and older adults both adopt strategies that reduce *MoS*_*L*_ [21, 23], reducing their base-of-support widths and lateral CoM motions. However, older adults have exhibited either smaller [23] or larger [19] average *MoS*_*L*_ relative to younger adults during narrow-base walking. Older adults’ larger (relative) mean *MoS*_*L*_ on narrow paths has been interpreted to indicate a preference for maintaining “safety” [19]. Conversely, smaller *MoS*_*L*_ may arise from their relatively larger, faster and more variable mediolateral CoM excursions [21, 24]. Thus, it is not clear how older adults alter *MoS*_*L*_ to remain on narrow paths or how these adjustments affect their lateral balance.

Here, younger and older adults traversed systematically-narrowing walking paths. This prompted participants to continually adjust their walking, allowing us to characterize how their mediolateral balance changed. We expected participants would exhibit gradually smaller and less variable *MoS*_*L*_ as paths narrowed [23]. We expected this would yield progressively larger *PoI*_*L*_, indicating greater likelihood of dropping below the *MoS*_*L*_ = 0 threshold on any given step. We expected older adults would exhibit consistently larger [19] but also more variable *MoS*_*L*_ [23] than younger adults. Thus, despite their increased *mean MoS*_*L*_, we expected older adults’ increased *MoS*_*L*_ *variability* would produce consistently larger *PoI*_*L*_, consistent with their greater risk of becoming laterally unstable. Here, we allowed participants to leave the narrowing path at any time [25]. We therefore also quantified how groups differed when leaving the narrowing path.

## METHODS

### Participants

We analyzed data from 20 younger (YH: 9M/11F, 21.7±2.6 years) and 18 older (OH: 5M/13F, 71.6±6.0 years) healthy adults described in [25, 26]. All participants provided signed informed consent, approved by Pennsylvania State University’s IRB. All participants scored ≥ 24/30 on the Mini-Mental State Exam [27] and had no injuries, prior surgeries or other walking impairments. Assessments included Timed Up and Go (TUG; [28]), Four Square Step Test (FSST; [29]), and abbreviated Iconographical-Falls Efficacy Scale (I-FES; [30]).

### Protocol and Data Collection

As detailed in [25], participants walked five 4-minute trials on a 1.2m wide treadmill (Motek M-Gait, Amsterdam, Netherlands). Each trial was 200m long and included eight bouts of two parallel paths, 25m long, projected onto the left and right treadmill belts (Fig. 1A-B). One path gradually narrowed from a width of *W*_*P*_ = 0.45m to *W*_*P*_ = 0.05m wide. The adjacent path remained fixed at 0.45m wide. Every 25m, the narrowing and fixed-width paths switched sides (Fig. 1C) and a new bout started.

**Figure 1:**
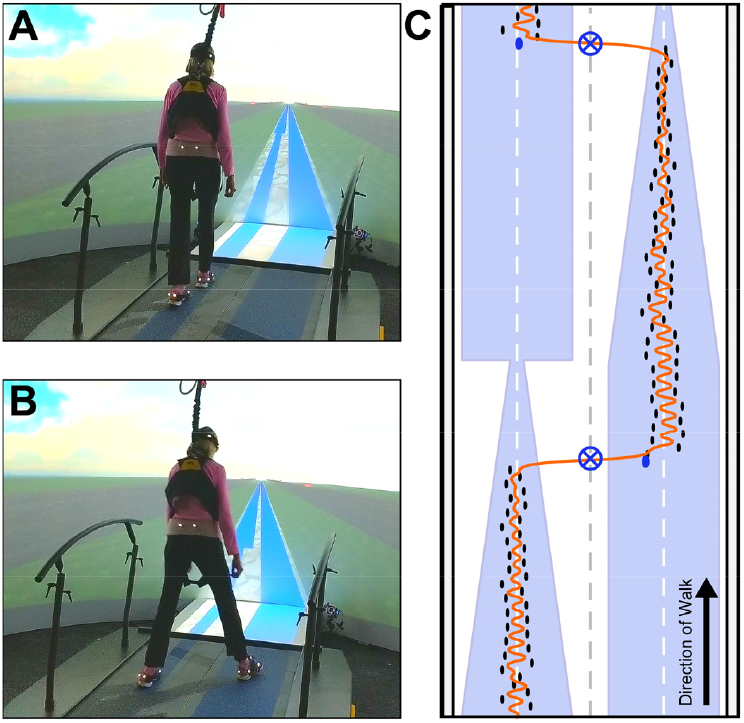
**(A)**: Participants walked on a 25m-long, gradually-narrowing path projected onto one belt of a split-belt treadmill surface. The width of the narrowing path gradually decreased from 0.45 to 0.05 m. A second “easier” path was projected onto the adjacent treadmill belt. This adjacent path remained at a fixed width of 0.45m. **(B)**: Participants were instructed to switch onto this adjacent path when they could no longer safely or successfully traverse the narrowing path. **(C)**: Each walking trial consisted of eight such walking bouts. Narrowing vs. fixed-width paths switched sides of the treadmill every 25m, initiating each new walking bout. Participants walked within the path bounds (black markers indicate foot placements) approximately along the path center (dashed white lines). Path switches (blue crosshairs) occurred when the mediolateral center-of-mass position (orange) crossed the centerline of the treadmill (dashed grey line). Transition steps occurred as the first step onto the fixed-width path for each walking bout (blue markers).

Participants were instructed to stay on each narrowing path as long as possible (Fig. 1A). Participants chose when to switch to the adjacent path (Fig. 1B). After each trial, participants completed 1 minute of steady walking and rested ≥2 minutes. For all trials, we fixed the treadmill speed at 0.75m/s, as older adults select similar speeds during narrow-base gait [21] and keeping speed fixed prevented confounds from different self-selected speeds.

Participants wore 16 retroreflective markers: four each on the head, pelvis, and each shoe [25]. A 10-camera Vicon system (Oxford Metrics, Oxford, UK) recorded marker trajectories at 120 Hz. We post-processed raw marker data in Vicon Nexus and Motek D-Flow software, then exported to MATLAB (MathWorks, Natick, MA). Marker trajectories were low-pass filtered (4^th^-order Butterworth, cutoff: 10Hz) and interpolated to 600Hz to compute accurate stepping events [31].

### Lateral Margins of Stability

We defined steps at consecutive heel strikes [32]. We analyzed only steps taken on the narrowing paths. These included all steps from the beginning of each bout until the second-to-last step before participants switched paths.

Lateral (*z*-direction) motion of the pelvic centroid approximated mediolateral center-of-mass (CoM) state (*z, ż*) at each time instant (Fig. 2A). The *z*-position of the leading foot’s 5^th^ metatarsal-phalangeal joint marker approximated the lateral base-of-support boundary (*u*_*max*_). The extrapolated CoM position [10] was then (*z* + *ż/ω*_0_), where 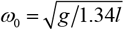, *g* = 9.81 m/s^2^, and *l* was vertical distance from the lateral malleolus to greater trochanter [10, 13]. The continuous-time (600Hz) lateral *Margin of Stability* was then [10]:

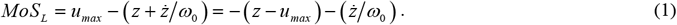

**Figure 2:**
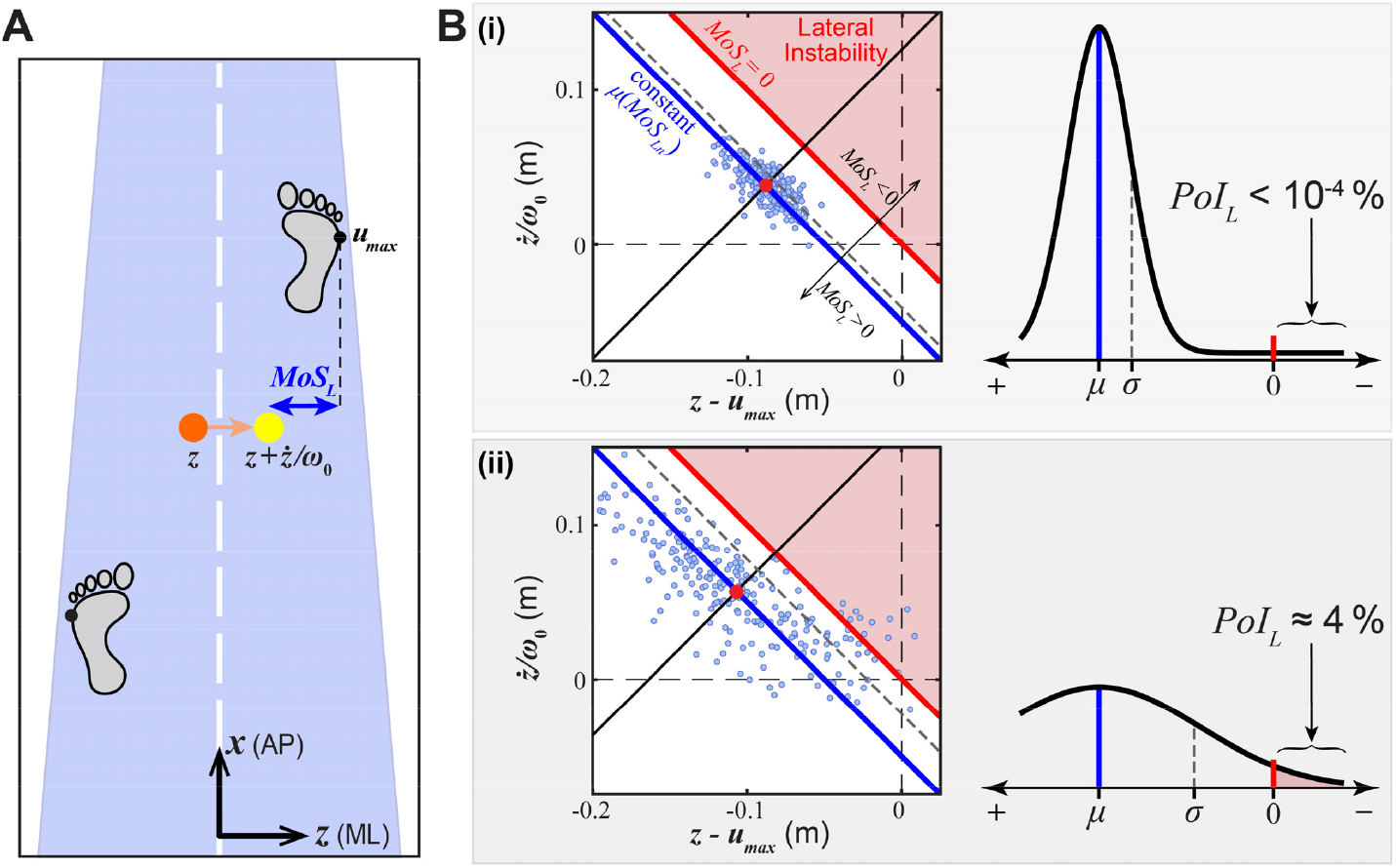
**(A)**: Schematic showing variables extracted at each step. The 5^th^ metatarsal-phalangeal joint (5MTP) marker of the leading foot approximated each foot’s lateral base-of-support boundary (*u*_*max*_). The pelvic centroid approximated the lateral position of the center-of mass (*z*). This was adjusted by the normalized velocity (*ż/ω*_0_) to define the lateral position of the extrapolated center-of mass (*z* + *ż/ω*_0_) [10]. Lateral Margins of Stability (*MoS*_*L*_) were then calculated as [(*u*_*max*_) – (*z* + *ż/ω*_0_)] (Eq. 1). **(B)**: Schematic showing variables relevant to calculating the lateral Probability of Instability (*PoI*_*L*_) for two cases: (i) low step-to-step variability and (ii) high step-to-step variability. For each case, **left panels** show minimum *MoS*_*Ln*_ values for individual steps (blue markers) plotted in the [*z*− *u*_*max*_, *ż/ω*_0_] plane. Solid red diagonal lines indicate lines of constant *MoS*_*L*_ = 0. Above these lines is the “Lateral Instability” region, where all *MoS*_*L*_ < 0 and thus fail to meet Hof’s dynamic stability condition [10]. For each distribution of steps (*MoS*_*Ln*_), solid blue diagonal lines indicate lines of constant average *μ*(*MoS*_*Ln*_) and dashed gray diagonal lines indicate each corresponding [*μ*(*MoS*_*Ln*_) − *σ*(*MoS*_*Ln*_)]. Note that cases (i) and (ii) exhibit nearly identical *μ*(*MoS*_*Ln*_), but case (ii) exhibits a larger *σ*(*MoS*_*Ln*_) and hence more steps taken in the lateral instability region. For each case, **right panels** show the probability distributions fitted to all *MoS*_*Ln*_ shown in each corresponding left panel. The lateral Probability of Instability (*PoI*_*L*_; Eq. (2)) for each distribution is the area under each curve in the Lateral Instability (i.e., *MoS*_*L*_ < 0) region. Thus, despite nearly identical *μ*(*MoS*_*Ln*_), case (ii) results in a much larger *PoI*_*L*_, due to its larger *σ*(*MoS*_*Ln*_).

For each *n*^*th*^ step taken on the narrowing path, we then extracted the minimum *MoS*_*L*_ within that step, *MoS*_*Ln*_, its corresponding (*z*−*u*_*max*_)_*n*_ and (*ż/ω*_*0*_)_*n*_, and the path width at that step, (*W*_*P*_)_*n*_. For each bout performed by each participant, this yielded time series of each of these variables for all steps taken during that bout.

### MoS_Ln_ Trends as Paths Narrowed

We first pooled time series of *MoS*_*Ln*_ for each group: YH and OH. As in [25, 26], we then divided the pooled group data into overlapping bins that spanned the range of *W*_*P*_. Each bin contained all *MoS*_*Ln*_ within a 0.075m range of *W*_*P*_, with new bins assigned every 0.005m of *W*_*P*_. Each bin was assigned a designated *W*_*P*_, taken as the mid-point of possible *W*_*P*_ for that bin [25].

As in [26], within each bin, we then extracted 50 data sets by sub-sampling a random 95% of the *MoS*_*Ln*_ in that bin. For each sub-sample, we computed one estimate (each) of the mean, *μ*(*MoS*_*Ln*_), and standard deviation, *σ*(*MoS*_*Ln*_), of all *MoS*_*Ln*_ in that bin. We then computed the overall means and between-sub-sample ±SD for all *μ*(*MoS*_*Ln*_) and *σ*(*MoS*_*Ln*_) estimates in each bin, each as a function of the narrowing path width, *W*_*P*_.

### Lateral Probability of Instability

Each step’s *MoS*_*Ln*_ can be visualized by plotting [(*z*−*u*_*max*_)_*n*_, (*ż/ω*_*0*_)_*n*_] in the [*z*−*u*_*max*_, *ż/ω*_*0*_] plane (Fig. 2B) [13]. In this plane, diagonal lines of slope −1 form manifolds of constant *MoS*_*L*_: i.e., all [(*z*−*u*_*max*_), (*ż/ω*_*0*_)] lying on any one such diagonal line have the exact same *MoS*_*L*_. For any sequence of *MoS*_*Ln*_, one such manifold occurs at *μ*(*MoS*_*Ln*_) (blue diagonal lines; Fig. 2B). A parallel manifold exists for *MoS*_*L*_ = 0 (red diagonal lines; Fig. 2B), which defines the threshold for “Lateral Instability” in the [*z*−*u*_*max*_, *ż/ω*_*0*_] plane [13]: any *MoS*_*Ln*_ beyond this threshold has *MoS*_*L*_ < 0 and fails Hof’s dynamic stability condition [10].

As evident from Fig. 2B, *μ*(*MoS*_*Ln*_) cannot predict the likelihood a person might experience a laterally unstable (i.e., *MoS*_*Ln*_ < 0) step. We therefore proposed the lateral *Probability of Instability* (*PoI*_*L*_) [13], which is the percent likelihood any given step will have *MoS*_*Ln*_ < 0. We estimated *PoI*_*L*_ as the probability (P) [13]:

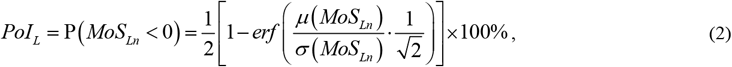

where *erf*(○) is the error function ([33]; https://dlmf.nist.gov/). Eq. (2) computes the integral of the probability density function over *MoS*_*L*_ < 0 for an assumed normal distribution of *MoS*_*Ln*_ having mean *μ*(*MoS*_*Ln*_) and standard deviation *σ*(*MoS*_*Ln*_) (Fig. 3B).

**Figure 3:**
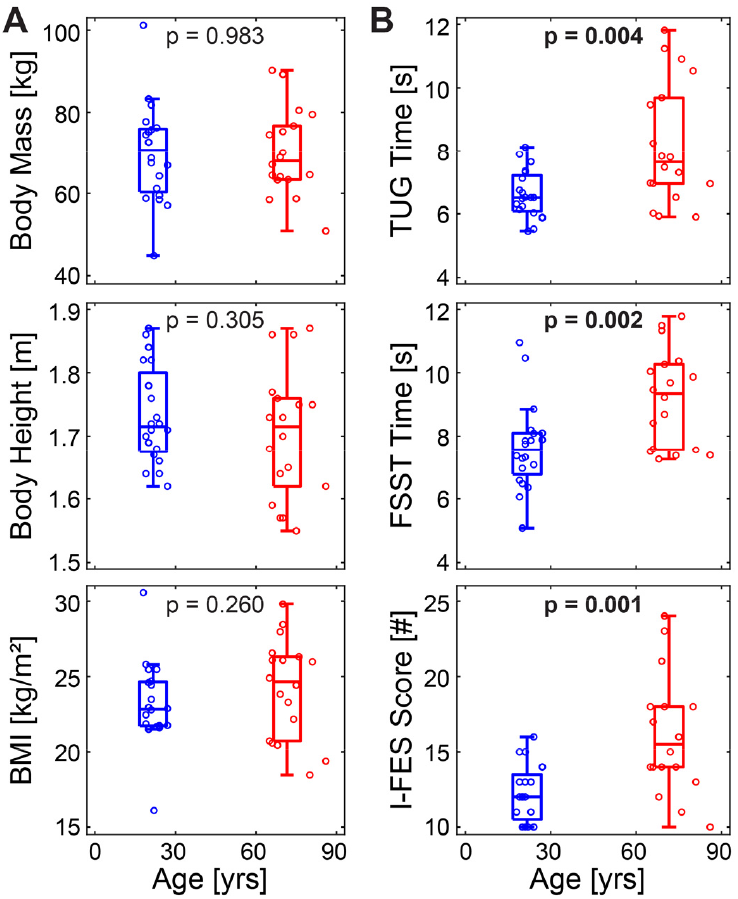
**A)** Demographics (body mass, height, and BMI) for each participant and group, plotted versus participants’ ages. Individual markers represent individual participants in each group (YH blue; OH red). Box plots are plotted at the mean age for each group. Each box plot shows the median, 1^st^ and 3^rd^ quartiles, and whiskers extending to 1.5× the inter-quartile range for that group. Groups did not differ in any of these characteristics (all p ≥ 0.26). **B)** Baseline assessments (TUG, FSST, and I-FES) for each participant and group, plotted in the same manner as in **(A)**. Compared to YH adults, OH adults exhibited significantly slower (more impaired) physical performance (TUG; p = 0.004, and FSST; p = 0.002), and were significantly more fearful of falling (I-FES; p = 0.001).

Here, we used the *μ*(*MoS*_*Ln*_) and *σ*(*MoS*_*Ln*_) calculated from each sub-sample within each bin in Eq. (2) to compute corresponding estimates of *PoI*_*L*_ for each sub-sample. We then computed the overall means and between-sub-sample ±SD for all *PoI*_*L*_ estimates from each bin, again as a function of *W*_*P*_.

### Path Switching

We recorded path-switches when the pelvic centroid crossed the treadmill midline (Fig. 1C) and we extracted the path widths at which people switched: (*W*_*P*_)_*switch*_. For each participant, we selected the appropriate group-wise bin closest to their mean (*W*_*P*_)_*switch*_. We used that bin’s average {*μ*(*MoS*_*Ln*_)_*switch*_, *σ*(*MoS*_*Ln*_)_*switch*_, (*PoI*_*L*_)_*switch*_} to characterize that participant’s lateral balance when they switched paths. We ran Mann-Whitney U tests (Minitab, Inc., State College, PA) to assess group (YH vs. OH) differences in each measure.

## RESULTS

YH and OH participants exhibited similar physical characteristics including body mass, height, and BMI (all p ≥ 0.26; Fig. 3A). However, OH exhibited slower TUG and FSST times (indicating impaired balance) and greater Icon-FES scores (indicating greater concern about falling) relative to YH (all p ≤ 0.004; Fig. 3B).

Figure 4 shows stepping data for a single walking bout for a representative participant. As the path narrows, step widths (*u*_*max*_) tend to narrow, lateral CoM (*z*) fluctuations decrease, and *MoS*_*Ln*_ tend to get smaller.

**Figure 4:**
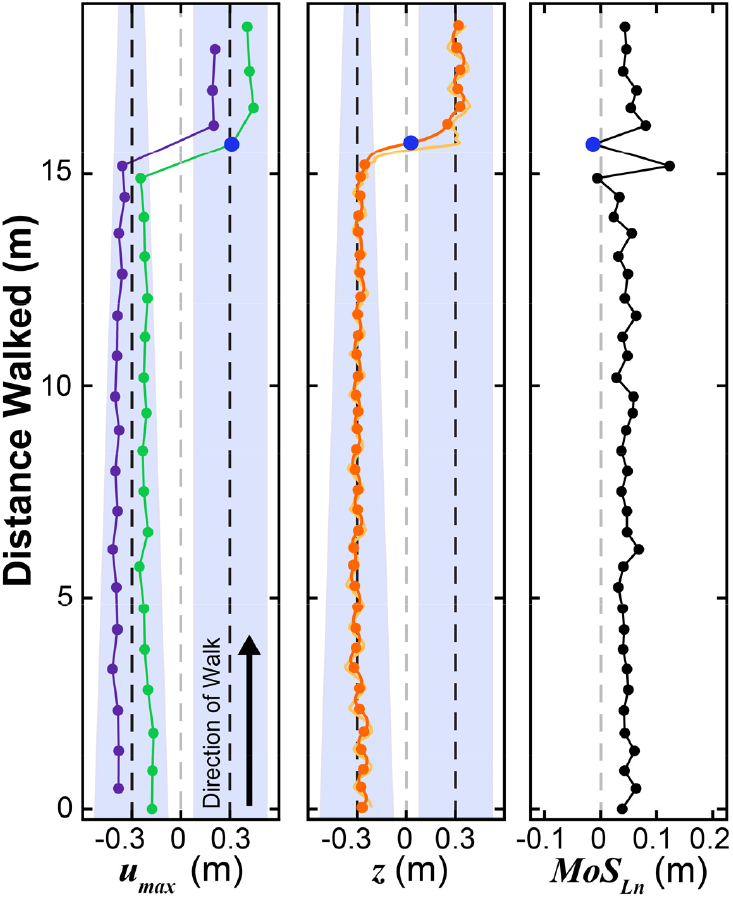
Example time series of relevant mediolateral balance variables at each step from a typical participant during a single walking bout. **Left:** lateral base-of-support boundary (*u*_*max*_) of left (purple) and right (green) foot placements. **Middle:** mediolateral CoM (*z*) (orange) and extrapolated CoM (*z* + *ż/ω*_0_) (light orange). **Right:** Corresponding minimum stability margins (*MoS*_*Ln*_) for each step. In each plot, a solid blue marker indicates the respective value at the transition step. In *u*_*max*_ and *z* plots, black dashed lines indicate path midlines and the grey dashed line indicates the treadmill midline. In the *MoS*_*Ln*_ plot, the grey dashed line indicates *MoS*_*L*_ = 0.

Across participants, as paths narrowed, all participants exhibited progressively smaller *z−u*_*max*_ distances and slower *ż/ω*_0_, and progressively smaller *MoS*_*Ln*_ (Fig. 5A). These resulted in altered group-wise distributions of [(*z−u*_*max*_)_*n*_, (*ż/*ω_0_)_*n*_], such that both groups’ *μ*(*MoS*_*Ln*_) diagonals tended to approach the *MoS*_*L*_ = 0 diagonal, contributing to increased *PoI*_*L*_ as paths narrowed (Fig. 5B). Interestingly, as people narrowed their step widths as the paths narrowed (i.e., mean (*z−u*_*max*_)_*n*_ shifted to the right), they also reduced their lateral CoM velocities (i.e., mean (*ż/*ω_0_)_*n*_ shifted downward). These reduced CoM velocities helped then to offset the narrower foot placements to keep their *μ*(*MoS*_*n*_) farther from the *MoS*_*L*_ = 0 diagonal (Fig. 5B).

**Figure 5:**
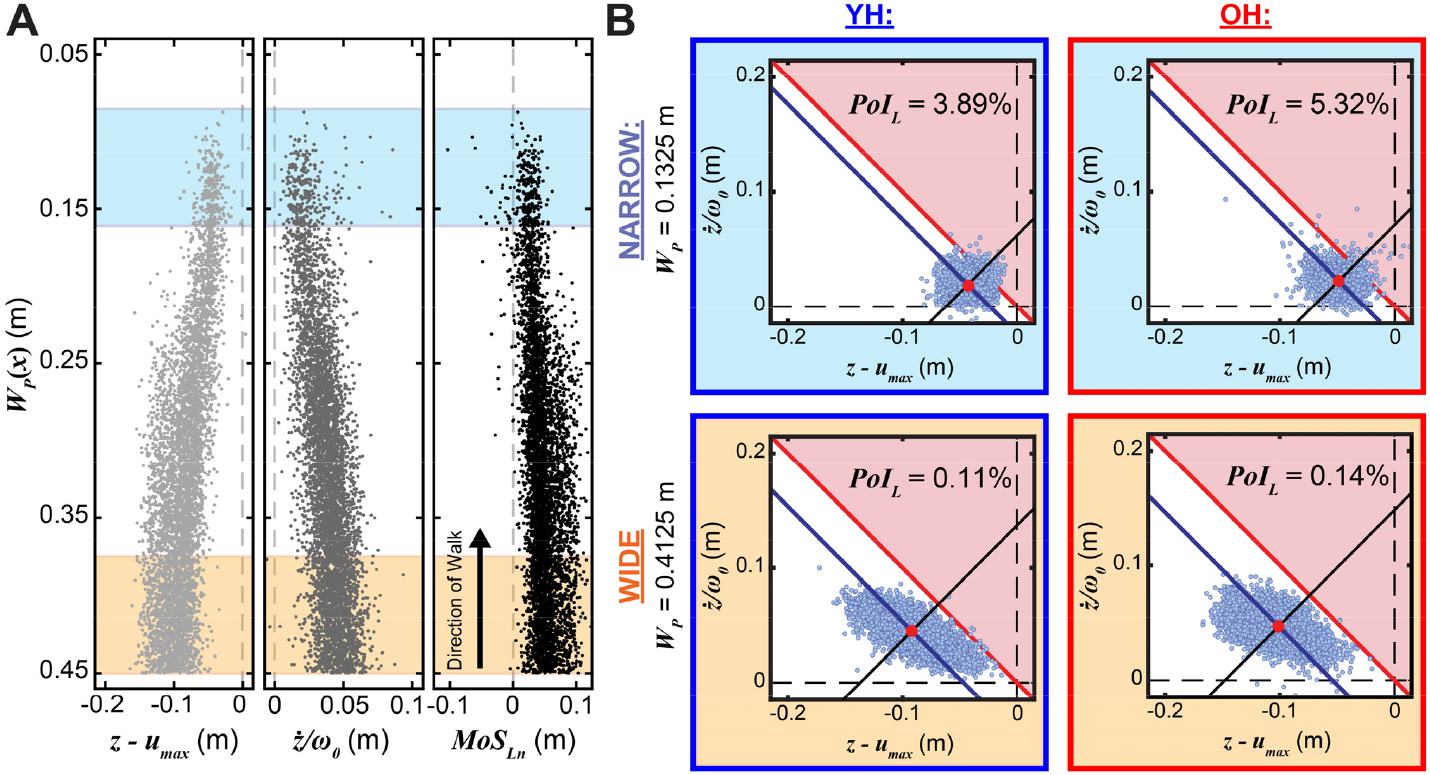
**(A)**: Adjusted mediolateral center-of mass positions (*z− u*_*max*_; left), normalized velocities (*ż*/ω_0_; middle), and lateral stability margins (*MoS*_*Ln*_; right) for all steps from five OH participants plotted vs. path widths, *W*_*P*_(*x*). Each marker represents a single step. On each plot, vertical grey dashed lines represent values equal to 0. Orange horizontal bands denote the first bin (see Methods) centered at *W*_*P*_ = 0.4125m containing all steps taken on the widest sections of the paths between *W*_*P*_ = 0.450m and *W*_*P*_ = 0.375m. Blue horizontal bands denote bins centered at *W*_*P*_ = 0.1325m containing all steps taken on paths between *W*_*P*_ = 0.170 m and *W*_*P*_ = 0.095m). (**B):** Group-wise mediolateral *MoS*_*Ln*_ coordinates plotted in the in the [*z− u*_*max*_, *ż*/ω_0_] plane (see Fig. 2B) for all steps for YH (left sub-plots) and OH (right sub-plots) groups for both the wide path width (*W*_*P*_ = 0.4125m; bottom sub-plots) and narrow path width (*W*_*P*_ = 0.1325m; top sub-plots) shown in (A). In each sub-plot, single markers indicate individual steps and solid blue diagonal lines indicate lines of constant *μ*(*MoS*_*Ln*_), as in Fig. 2B. At narrower path widths (*W*_*P*_ = 0.1325m), these distributions become more isotropic, with *μ*(*MoS*_*Ln*_) diagonals closer to the lateral instability region and substantially larger *PoI*_*L*_. OH participants’ *PoI*_*L*_ (5.32%) is 37% larger than that of YH (3.89%).

As paths narrowed, both groups progressively reduced their *μ*(*MoS*_*Ln*_), while OH exhibited consistently larger *μ*(*MoS*_*Ln*_) relative to YH (Fig. 6A). Initially, both groups steadily reduced their *σ*(*MoS*_*Ln*_) as paths narrowed, but then increased their *σ*(*MoS*_*Ln*_) as paths narrowed further (Fig. 6B). OH exhibited consistently greater *σ*(*MoS*_*Ln*_) relative to YH (Fig. 6B). Both groups exhibited *PoI*_*L*_ that increased sharply as paths narrowed (Fig. 6C). OH exhibited somewhat larger *PoI*_*L*_ relative to YH on narrower path sections (Fig. 6C).

**Figure 6:**
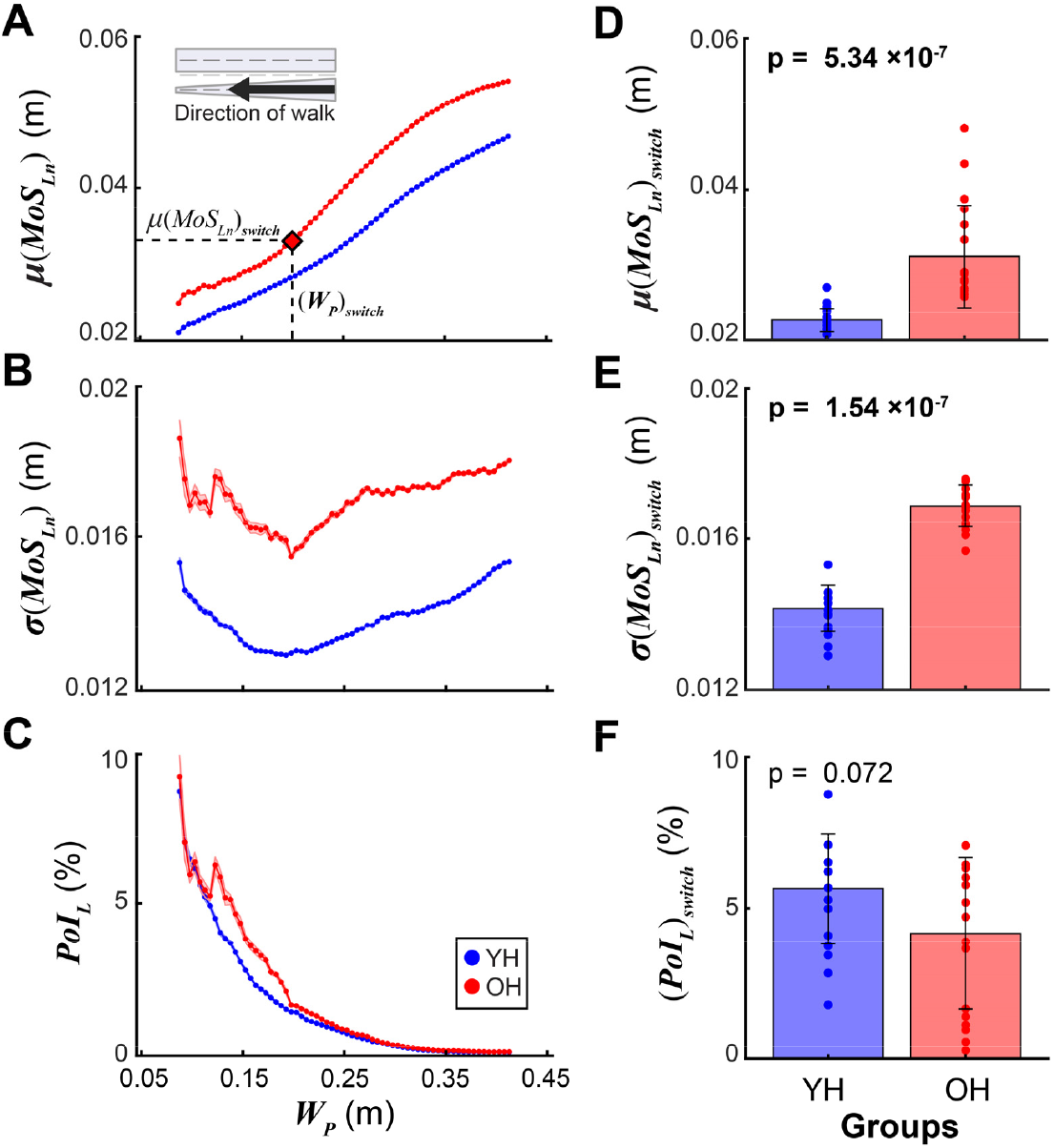
Lateral margin of stability **(A)** means *μ*(*MoS*_*Ln*_), **(B)** variance *σ*(*MoS*_*Ln*_), and **(C)** lateral probability of instability (*PoI*_*L*_) across bins of decreasing path widths (*W*_*P*_). *μ*(*MoS*_*Ln*_), *σ*(*MoS*_*Ln*_), and *PoI*_*L*_ were each averaged across all sub-samples within each bin (see Fig. 5). Markers represent bin averages. Shaded regions (small, but present) represent between sub-samples ±1SD bands. The path width bin at which each participant switched paths, (*W*_*P*_)_*switch*_, was mapped onto each of these plots to characterize their balance at path switching, for example, as shown in **(A)** for *μ*(*MoS*_*Ln*_)_*switch*_. Compared to YH adults, OH exhibited (**D**) larger *μ*(*MoS*_*Ln*_)_*switch*_ (**p = 5**.**34×10**^**-7**^) and (**E**) larger *σ*(*MoS*_*Ln*_)_*switch*_ (**p = 1**.**54×10**^**-7**^), but (**F**) similar (*PoI*_*L*_)_*switch*_ (p = 0.072).

Most OH adults switched off the narrowing path sooner than YH, at wider *W*_*P*_ (p = 0.022; data published in [25, 26]). When participants switched paths (i.e., at (*W*_*P*_)_*switch*_), OH participants exhibited significantly larger *μ*(*MoS*_*Ln*_)_*switch*_ (Fig. 6D) and *σ*(*MoS*_*Ln*_)_*switch*_ (Fig. 6E), but similar (*PoI*_*L*_)_*switch*_ (Fig. 6F) relative to YH.

## DISCUSSION

During daily walking, people must adjust their balance while maneuvering in constrained environments [2-4], including narrower paths. Older adults may be less able to adapt to such circumstances [6, 7, 21]. Walking on narrow paths amplifies age-related balance impairments [18-24]. Unlike those prior studies, here participants walked down *continuously*-narrowing paths (Fig. 1) that progressively challenged their balance. We quantified their CoM dynamics (Fig. 5) that determine instability risk (Fig. 6).

As paths narrowed, participants gradually narrowed their steps toward their midline [26] (decreasing *z−u*_*max*_; Fig. 5A). As they did, they also reduced their lateral CoM velocities (decreasing *ż/*ω_0_; Fig. 5A). Hence, they exploited the redundancy intrinsic in the [(*z−u*_*max*_), (*ż/*ω_0_)] plane (Fig. 2B; [13]): decreases in *z−u*_*max*_ were partly compensated by decreases in *ż/*ω_0_, which limited deviations from their starting constant *μ*(*MoS*_*Ln*_) diagonal (Fig. 5B). Such compensations likely underlie prior findings suggesting people try to maintain *μ*(*MoS*_*Ln*_) as they walk [34, 35].

All participants exhibited progressively smaller *μ*(*MoS*_*Ln*_) as they walked down the narrowing paths (Fig. 6A). Participants exhibited largest *σ*(*MoS*_*Ln*_) on both the widest path sections that afforded the most stepping options, and the narrowest path sections that required the greatest stepping accuracy (Fig. 6B). However, despite reducing *ż/*ω_0_ to try to limit deviations from their starting *μ*(*MoS*_*Ln*_) (Fig. 5B), these narrowing paths eventually forced participants’ CoM states to draw closer to the *MoS*_*L*_ = 0 threshold (Figs. 5B, 6A). This, along with their increased *σ*(*MoS*_*Ln*_) (Fig. 6B) contributed to their dramatically increased *PoI*_*L*_ as paths narrowed (Fig. 6C).

Despite older participants (OH) exhibiting clear balance impairments on standard clinical tests (Fig. 1B), they consistently walked with *larger* (not smaller) *μ*(*MoS*_*Ln*_) than YH (Fig. 6A). Though consistent with many similar findings [12-16, 19], interpreting larger *μ*(*MoS*_*Ln*_) as indicating that OH participants were *more* (not less) stable than YH would be both highly counterintuitive and contrary to Hof’s proposition [10, 11]. This is because *μ*(*MoS*_*Ln*_) ignores the larger *variability* OH exhibited (Fig. 6B). Here, OH adults simultaneously remained *farther* (on average) from the *MoS*_*L*_ = 0 lateral stability threshold (Figs. 5B, 6A), yet *more* likely to take unstable (*MoS*_*Ln*_ < 0) steps on narrower path regions (Fig. 6C). Aging elicits greater neurophysiological noise [36] that contributes to increased gait variability [7, 37]. Older adults must contend with this variability to maintain balance. Here, OH adults took wider steps [26], that helped increase their *μ*(*MoS*_*Ln*_) (Fig. 6A) [38]. This directly mitigated their balance risk. If OH adults had maintained the smaller *μ*(*MoS*_*Ln*_) of YH (Fig. 6A), but retained their increased *σ*(*MoS*_*Ln*_) (Fig. 6B), they would have had much higher *PoI*_*L*_ (Fig. 6C) than they did. Our findings thus explain precisely *why* wider steps help OH maintain balance [39] and resolves the paradox [13] of misinterpreting *μ*(*MoS*_*Ln*_). We assert this is because *PoI*_*L*_ (Fig. 6C) directly measures statistical *risk* of instability, consistent with Hof’s proposition [10, 11], while *μ*(*MoS*_*Ln*_) does not.

We emphasize that while *μ*(*MoS*_*Ln*_) clearly does not predict *falling* risk (Fig. 2B; [13]), neither does *PoI*_*L*_. Instead, *PoI*_*L*_ predicts one’s likelihood of experiencing *MoS*_*Ln*_ < 0 on a given step. Experiencing *MoS*_*Ln*_ < 0 does not automatically necessitate a fall (e.g., here, no one fell despite numerous steps below that threshold; Fig. 5). Hence, *MoS*_*Ln*_ is not a useful measure of stability in a deterministic sense. Rather, *MoS*_*Ln*_ < 0 indicates only that alternative strategies not available to an inverted pendulum [10] must be used to restore balance. Conversely, *PoI*_*L*_ quantifies how people balance their mean-variance trade-off [40] between *μ*(*MoS*_*Ln*_) and *σ*(*MoS*_*Ln*_) to avoid potential falls. Our results indicate that people attempt to keep *PoI*_*L*_ below some personally-acceptable level, depending on their *risk-sensitivity* [40].

Older adults left the narrowing paths sooner at wider *W*_*P*_ than young adults [25, 26], as mediated by their lower self-perceived balance abilities [25]. Here, just prior to transitioning, older adults exhibited larger, more variable *MoS*_*Ln*_ and slightly smaller *PoI*_*L*_ than younger (Fig. 6D-F). Thus, these older adults, despite lower physical and self-perceived balance capabilities (Fig. 1B; [25]), left the narrowing path while still exhibiting similar-to-lower *PoI*_*L*_ as our young adults (Fig. 6F). This suggests OH decided to switch paths at path widths where they took on comparable levels of instability risk as YH.

One limitation of our paradigm was that participants chose when to switch off the narrowing paths, which we designed to be nearly impossible to walk on at their narrowest (5cm). We did this intentionally to quantify participants’ path-switching decisions [25]. However, different participants switching at different *W*_*P*_ resulted in fewer data points in each group-wise bin as the paths narrowed, and more so for older adults.

This study revealed for the first time how older adults adapt mediolateral balance adjustments to real-time changes in path constraints. Older adults exhibited degraded balance (Figs. 1B). However, they used multiple strategies to compensate. Specifically, during steady walking, they mitigated their increased *σ*(*MoS*_*Ln*_) (Fig. 6B) by taking wider steps to walk with increased *μ*(*MoS*_*Ln*_) (Fig. 6A), yielding *PoI*_*L*_ more comparable to those of younger participants (Fig. 6C). They also left the narrowing paths sooner [25], but again at comparable levels of instability risk (Fig. 6F). Our results directly challenge the predominant, yet problematic interpretation that larger *μ*(*MoS*_*Ln*_) indicate “greater stability” [12]. Conversely, despite exhibiting consistently *larger* (supposedly “more stable”) average *MoS*_*L*_ than younger (Fig. 6A), *PoI*_*L*_ [13] (Fig. 6C) correctly characterized older adults’ instability risk. Furthermore, the adaptive strategies uncovered using these more nuanced methods suggest potential targets for future interventions seeking to prevent falls in older adults.

## ACKNOWLEDGEMENTS

The authors thank Anna C. Render and David M. Desmet for their contributions and technical support throughout data collections. This work was supported by NIH grant 1-R01-AG049735 to JBD & JPC.

## Notes

**CONFLICT OF INTEREST**: The authors declare that there is no conflict of interest associated with this work.

### Competing Interest Statement

The authors have declared no competing interest.

